# Growth hormone transgenesis disrupts immune function in muscle of coho salmon (*Oncorhynchus kisutch*) impacting cross-talk with growth systems

**DOI:** 10.1101/210104

**Authors:** Abdullah Alzaid, Jin-Hyoung Kim, Robert H. Devlin, Samuel A.M. Martin, Daniel J. Macqueen

**Affiliations:** School of Biological Sciences, University of Aberdeen, Aberdeen, AB24 2TZ, UK.; Fisheries and Oceans Canada, West Vancouver, British Columbia, V7V 1N6, Canada.; Current address: Korea Polar Research Institute (KOPRI), Yeonsu-gu, Incheon 21990, Korea.

**Author notes:** Corresponding authors: D.J.M. S.A.M.M.

**Keywords:** Growth, immunity, growth-immune cross-talk, skeletal muscle, growth hormone, insulin-like growth factor, transgenesis, coho salmon

## Abstract

The suppression of growth during infection should facilitate resource allocation towards effective immune function. Work supporting this hypothesis has been recently reported in teleosts, demonstrating immune-responsive regulation of the insulin-like growth factor (IGF) system - a key endocrine growth pathway that acts downstream of growth hormone (GH). Skeletal muscle is the main target for growth and energetic storage in fish, yet little is known about how growth is regulated in this tissue during an immune response. We addressed this knowledge gap by characterizing muscle immune responses in size-matched coho salmon *(Oncorhynchus kisutch)* achieving different growth rates. We compared a wild-type strain with two GH transgenic groups achieving either maximal or highly-suppressed growth – an experimental design that separates GH’s direct effects from its influence on growth rate. Fish were sampled 30h post-injection with PBS (control) or mimics of bacterial (peptidoglycan) or viral (Poly:IC) infection. We quantified the mRNA level expression of genes from the GH, GH receptor (GHR), IGF hormone, IGF1 receptor (IGF-1R) and IGF binding protein (IGFBP) families, along with marker genes for muscle growth and host defence genes involved in inflammatory or antiviral responses. We provide strong evidence for dampened immunity in the GH transgenics compared to wild-type animals. Strikingly, the muscle of GH transgenics achieving rapid growth showed no detectable antiviral response, coupled with evidence of a constitutive inflammatory state. GH and IGF system gene expression was also strongly altered by GH transgenesis and fast growth, both for baseline expression levels and responses to immune stimulation. Overall, our findings demonstrate that GH transgenesis disrupts normal immune function and growth-immune cross-talk in muscle, with implications for the health and welfare of farmed salmon.

## INTRODUCTION

The skeletal muscle is the most important target for growth investment and energy storage in teleost fish, representing more than half of body mass in salmonid species. This tissue is remobilised regularly during life, for example upon maturation or during fasting, with the resultant resources available for allocation into other physiological systems. The regulation of skeletal muscle mass represents a dynamic balance between protein synthesis and degradation pathways, controlled by growth hormone (GH) and insulin-like growth factors (IGFs) (Johnston et al., 2011; Fuentes et al. 2013). GH is the master growth regulator in all vertebrates, and acts on target tissues via its receptor GHR, or indirectly by stimulating hepatic production of IGFs (Fuentes et al., 2013). The IGF system comprises IGF-I and IGF-II hormones, which promote growth phenotypes through IGF-1R signalling pathways (Jones and Clemmons, 1995; Wood et al., 2005), including in skeletal muscle (Johnston et al., 2011). The action of IGFs is modulated in the extracellular environment by a family of IGF-binding proteins (IGFBPs) that influence the availability of these hormones to IGF-1R (Firth and Baxter, 2002). Many genes from the GH and IGF pathways have been expanded by genome duplication events in teleost evolutionary history (Ocampo Daza et al., 2011; Macqueen et al., 2013; Lappin et al., 2016; Alzaid et al., 2016a; Robertson et al., 2017), including a salmonid-specific event that occurred 88-103 Ma (Macqueen and Johnston, 2014).

Many past studies of teleosts have investigated the *in vivo* regulation of GH and IGF pathway genes under distinct nutritional states, highlighting a key role for these systems in the growth and remodelling of skeletal muscle. For example, Atlantic salmon *(Salmo salar* L.) fed after a period of fasting showed upregulation of *IGF-I* and *IGFBP-4,* whereas *IGF-II, IGF1R* and *IGFBP-2* were downregulated (Bower et al., 2008). Reciprocally, muscle *IGF-I* transcript levels were downregulated in Atlantic salmon (Breves et al., 2016), yellowtail *(Seriola quinqueradiata)* (Fukada et al., 2012) and tilapia *(Oreochromis mossambicus)* (Fox et al., 2010) by fasting, while *GHR* transcripts were induced by fasting in tilapia (Fox et al., 2010). Despite such gains in understanding, to the best of our knowledge, the regulation of GH and IGF pathway genes during an immune response in teleost skeletal muscle remains uncharacterized.

Past work provides evidence for cross-talk between the GH and IGF pathways and the immune system (e.g. Heemskerk et al. 1999; Yada, 2007; O’Connor et al., 2008; Smith, 2010; Franz et al., 2016). We recently reported that IGF signalling is repressed in rainbow trout *(Oncorhynchus mykiss)* in response to bacterial and viral infections, with a striking upregulation of IGFBP subtypes that restrict IGFs from IGF-1R, and potentially modify immune function (Alzaid et al., 2016a, 2016b). Interestingly, immune responsive genes from the IGF pathway showed a striking co-expression with pro-inflammatory cytokine and antiviral genes regulated by conserved immune signalling pathways (Alzaid et al., 2016b), supporting the hypothesis that mechanisms have evolved that limit growth investment as an intrinsic component of host defence.

What little is known about the regulation of GH and IGF pathway genes in teleost muscle following immune stimulation has come from *in vitro* work. Treatment of Atlantic salmon muscle cell cultures with the pro-inflammatory cytokine IL-1β resulted in up-regulation of an IGFBP-6 subtype, suggested to represses IGF signalling (Pooley et al., 2013; Heidari et al., 2015). However, in a separate study, treatment of differentiated Atlantic salmon myotubes with the same cytokine resulted in limited expression changes in IGF system genes, despite the verified presence of muscle fibre atrophy (Garcia de la serrana et al., 2017). Given the dearth of knowledge in this area, an important goal of the current study was to perform a comprehensive *in vivo* analysis of expression responses for GH and IGF pathway genes following immune stimulation in teleost skeletal muscle.

GH transgenesis offers an ideal model to address potential mechanisms of cross-talk between growth and immunity. Salmonid species overexpressing GH in a wild genetic background show enhanced appetite, feed intake and food conversion (e.g. Devlin et al. 2009; White et al. 2016), resulting in highly elevated growth rate (e.g. Devlin et al., 1994; 2015). As such, rapid growth requires a matched increase in energetic intake, but it is also possible for GH transgenic salmon to achieve suppressed growth (approaching wild-type) by provision of a wild-type ration (Rise et al. 2006; Raven et al., 2008). GH transgenic coho salmon on a restricted ration have the same plasma IGF-I levels and liver *IGF-I* mRNA expression as wild-type fish, despite highly increased plasma GH (Raven et al., 2008). This study system can be used to disentangle the impacts of GH from its influence on feed intake and growth rate. To date however, there are no published reports addressing the impact of GH transgenesis on the response of skeletal muscle to immune challenge.

The aim of the current study was to characterise gene expression regulation linking growth to immune function within the skeletal muscle of coho salmon, focussing on the GH and IGF systems. We contrasted transcriptional responses of genes from both pathways, in addition to selected markers of immune and muscle growth status, to immune stimulation in three experimental groups, comparing wild-type animals with a GH transgenic strain achieving either maximal or supressed growth by ration manipulation. Our findings reveal a disruption to immune function and the regulation of growth-immune cross-talk in muscle of GH transgenic animals, with implications for the health of rapidly growing fish strains used in aquaculture.

## MATERIALS AND METHODS

### Experimental design

Experiments on coho salmon were performed at Fisheries and Oceans Canada (DFO), West Vancouver, British Columbia, Canada. This facility is designed to prevent the escape of transgenic fish to the natural environment. All work was done in accordance with guidelines of the Canadian Council on Animal Care, under a permit (#12-017) from the DFO’s Pacific Regional Animal Care Committee. All studied fish were initially maintained under common garden conditions (4,000L tanks supplied with 10.5 ± 1 °C aerated well water, natural photoperiod, at a density <5 kg/m^3^) and fed a commercial diet (Skretting Ltd., Canada) twice daily at 09:00 and 15:00 (3% of body weight per day). Three experimental groups were generated after Oakes et al. 2007 and Raven et al. 2008: (i) 19-month-old wild-type animals fed to satiation throughout ontogeny (wild-type: ‘WT’), (ii) 6-month-old GH transgenic animals fed to satiation throughout ontogeny (transgenic full ration: ‘TF’), and (iii) 17-month-old GH transgenic animals fed to the WT satiety level throughout ontogeny (transgenic restricted ration: ‘TR’). Using fish of different ages was necessary to standardize the confounding effects of body size, owing to different growth rates among the groups. The WT group was the offspring of parents collected at the Chehalis River in British Columbia, Canada (Devlin et al., 2004). The GH transgenic strain (M77) was originally produced by microinjecting the GH gene construct of sockeye salmon *(O. nerka;* OnMTGH1) into the eggs of WT parents (Devlin et al., 1994, 2004), and subsequently maintained by crossing transgenic males to Chehalis River females at each generation to maintain a wild-type genetic background.

For each experimental group, 60 animals (size-matched, immature and of unknown sex; means±s.d. as follows: WT: 74.2±3.6 g, TF: 77.9±6.1 g, TR: 78.6±3.3 g) were marked by fin-clips and allocated into four separate 70L tanks prior to immune stimulation. The fish were then intraperitoneally injected with either: i) polyinosinic-polycytidylic acid (Poly:IC) at 200μg per 100g fish weight (24 fish per tank, per group), ii) peptidoglycan (PGN) at 200μg per 100g fish weight (24 fish per tank, per group) or iii) phosphate-buffered saline (PBS) (i.e. control, 24 fish per tank, per group). After treatment, fish were re-stocked into the 4,000L tanks and maintained under the common garden design described above. The concentrations of Poly:IC and PGN used were based on past studies (Kono and Sakai, 2001; Jensen et al. 2002; Lockhart et al. 2004; Kono et al. 2004). At the point of sampling, fish were killed by a lethal dose of tricaine methanesulphonate (200mg/L; Syndel Laboratories Ltd., Vancouver, BC, Canada; buffered in 400mg/L sodium bicarbonate) after prior sedation using Aquacalm (1mg/L; Syndel Laboratories Ltd., Vancouver, BC, Canada). For each group, 10 fish were randomly sampled 6h and 30h post-treatment, done at the exact same time of day for both time points. A panel of tissues, namely skeletal muscle, intestine, liver, head-kidney and spleen, were rapidly team sampled. For all tissues except skeletal muscle, samples were fixed in RNAlater™ (ThermoFisher Scientific) overnight at 4°C, and stored at −80°C. For skeletal muscle, the samples were split, with half fixed in RNAlater as described above, and the other half flash frozen on dry ice. For the current study, the skeletal muscle samples were shipped on dry ice to the School of Biological Sciences, University of Aberdeen, UK, where samples were stored at −70°C until analysis. Samples fixed in RNAlater were used for all molecular analyses described below *(n=5* fish per group, per treatment; 45 samples).

### Primers for quantitative PCR

Details of primers pairs for 47 quantitative PCR (qPCR) assays performed in the study are provided in Table S1, including citations to previously published primers. Coho salmon genes of interest were initially acquired using Atlantic salmon and rainbow trout orthologues acquired from the NCBI database as queries in BLASTn searches against two published coho salmon transcriptomes (Kim et al., 2016; Garcia de la Serrana et al. 2015) and a sequence capture dataset that included target genes from the GH and IGF systems for coho salmon (Lappin et al., 2016; Robertson et al., 2017). When the current paper was in preparation, a high-quality genome was released for coho salmon (NCBI accession; GCA_002021735.1). Hence, a larger pool of gene models became available, which were used to check all coho salmon sequences targeted by qPCR; where possible, we report coho-specific accession numbers for all gene targets (Table S1). For most IGF system genes, we found that published primers from Atlantic salmon (Macqueen et al., 2013) and rainbow trout (Alzaid et al., 2016a, 2016b) were conserved in coho salmon. New primer pairs were designed for *IGFBP-1A2* and *IGFBP-5B1* owing to significant mismatches between published primers and coho salmon. Salmonid-specific genes encoding GH are known for salmonids (previously named *GH1* and *GH2)* (e.g. McKay et al. 2004; Robertson et al. 2017) and both were identified in coho salmon (accession numbers in Table S1). We initially tested primers conserved across both GH duplicates and detected limited muscle transcript expression: since this primer pair binds both genes equally, we concluded that neither *GH* duplicate was sufficiently expressed to warrant design of additional primers. A past study identified salmonid-specific duplicates of GHR *(GHR1* and *GHR2),* including in coho salmon (Very et al. 2005), for which we designed new primer pairs that bind divergent regions among the duplicates (Table S1). Additional primers were used for marker genes known to be strongly upregulated by immune stimulation or to be directly involved in muscle growth and development (described in: Castro et al. 2015; Alzaid et al., 2016a, 2016b).

### Quantitative gene expression analyses

Total RNA extraction and reverse transcription was performed as described elsewhere (Alzaid et al., 2016b). Transcript levels of the target genes were measured with qPCR using an Mx3005P qPCR System with Brilliant III Ultra-Fast SYBR Green (Agilent Technologies). The efficiency of qPCR assays was calculated using LinRegPCR (Ruijter et al., 2009). Data analyses was performed in GenEx (MultiD Analyses AB) using the variance based algorithm NormFinder (Andersen et al., 2004) to test the suitability of five potential reference genes *(RpL4, RpS13, RpS29, ACTB* and *EF1A)* for data normalization; this approach identified *RpL13* and *ACTB* as the most stable pair of reference genes across all samples (combined s.d.: 0.14). The same genes were the most stably expressed considering variation in expression across treatments (Combined s.d.: 0.10) and fish groups (combined s.d.: 0.10), and were used to normalize the expression data for each experimental gene. Within GenEx, efficiency-corrected, normalized arbitrary transcript levels were placed on a relative scale that was quantitatively comparable across different genes.

### Statistics

Statistical analyses were performed in Minitab v.18 (Minitab Inc.). Differences in baseline gene transcript levels among fish groups for the control animals (PBS-injected) were identified using one-way ANOVA, with Tukey’s post-hoc test to reveal significant pair-wise differences among groups (i.e. WT-PBS vs. TR-PBS vs. TF-PBS). The effect of PGN and Poly:IC on gene expression was tested (separately for each immune mimic) using two-way ANOVA, including the effect of treatment, fish group and treatment-x-group interaction. When two-way ANOVA revealed a significant effect of treatment plus a significant treatment-x-group interaction, we used Tukey’s post-hoc test to: i) identify significant pair-wise differences within each fish group due to treatment (i.e. WT-control vs. WT-PGN or WT-Poly:IC; TR-control vs. TR-PGN or TR-Poly:IC; TF-control vs. TF-PGN or TF-Poly:IC) and ii) identify significant pair-wise differences in transcript levels among fish groups subjected to each immune treatment (i.e. WT-PGN vs. TR-PGN vs. TF-PGN; WT-Poly:IC vs. TR-Poly:IC vs TF-Poly:IC). For all parametric analyses, we tested whether the fitted model residuals conformed to assumptions of normality (Anderson-Darling test) and homoscedasticity (Levene’s test). Box-Cox transformations and if necessary a nonparametric test (Kruskal-Wallis) were employed when data failed to meet these assumptions.

## RESULTS

### GH transgenesis alters baseline expression of GH and IGF system genes in skeletal muscle

We first assayed the baseline mRNA levels of all tested GH and IGF pathway genes in the muscle of unstimulated control fish for the three experimental groups (Fig. 1; Table 1). For the mRNAs encoding hormones, *GH* was expressed at low levels in all groups, *IGF-I* expression was not different across the groups, while *IGF-II* expression was significantly elevated (by ~2.3 fold) in TR vs. WT (Fig. 1A; Table 1). *IGF-II* levels were substantially (~20 fold) higher than *IGF-I* in all groups (Table 1). Among the assayed receptors, *GHR1* transcript levels were significantly lower in TF vs. WT and TF vs. TR comparisons (by ~4.5 and 2.5 fold, respectively), whereas *GHR2* expression was not significantly different across groups (Fig. 1B; Table 1). Expression of *IGF1R-a2* was higher than other IGF1R family genes (i.e. *IGF1R-a1* and *IGF1R-b)* in all groups, and significantly higher in TF vs. WT, by ~2.4 fold (Fig. 1C; Table 1). The expression of four out of eleven expressed IGFBP family member genes differed significantly between the three groups (Table 1). No muscle expression was detected for *IGFBP-1B1,-1B2,-2B1,-2B2,-3B1,-3B2,-6A1* and *-6A2. IGFBP-1A2* transcript levels were significantly higher (by ~3.8 fold) in TF vs. WT, but were not significantly different comparing TF and TR, despite showing a trend of being higher in the former (Fig. 1D). Conversely, *IGFBP-3A1, IGFBP-5B1* and *IGFBP-6B2* were each most highly expressed in the TR group, and always significantly higher than WT (and significantly higher than TF for *IGFBP-5B1* only) (Fig. 1E-G; Table 1). For the tested markers of muscle growth status, most were not differentially expressed across groups, including *FBXO32* (Tacchi et al., 2010), which encodes an E3-ubiquitin ligase involved in structural protein turnover, *TNNI2* and *MYL1,* which encode sarcomere proteins and *MyoG,* a transcription factor for myogenic differentiation (Table 1). However, transcript levels of *MyoD1a* (Macqueen and Johnston, 2006), a transcription factor for myogenic determination and differentiation, were significantly elevated in TF vs. WT (Fig. 1H; Table 1).

**Figure 1.**
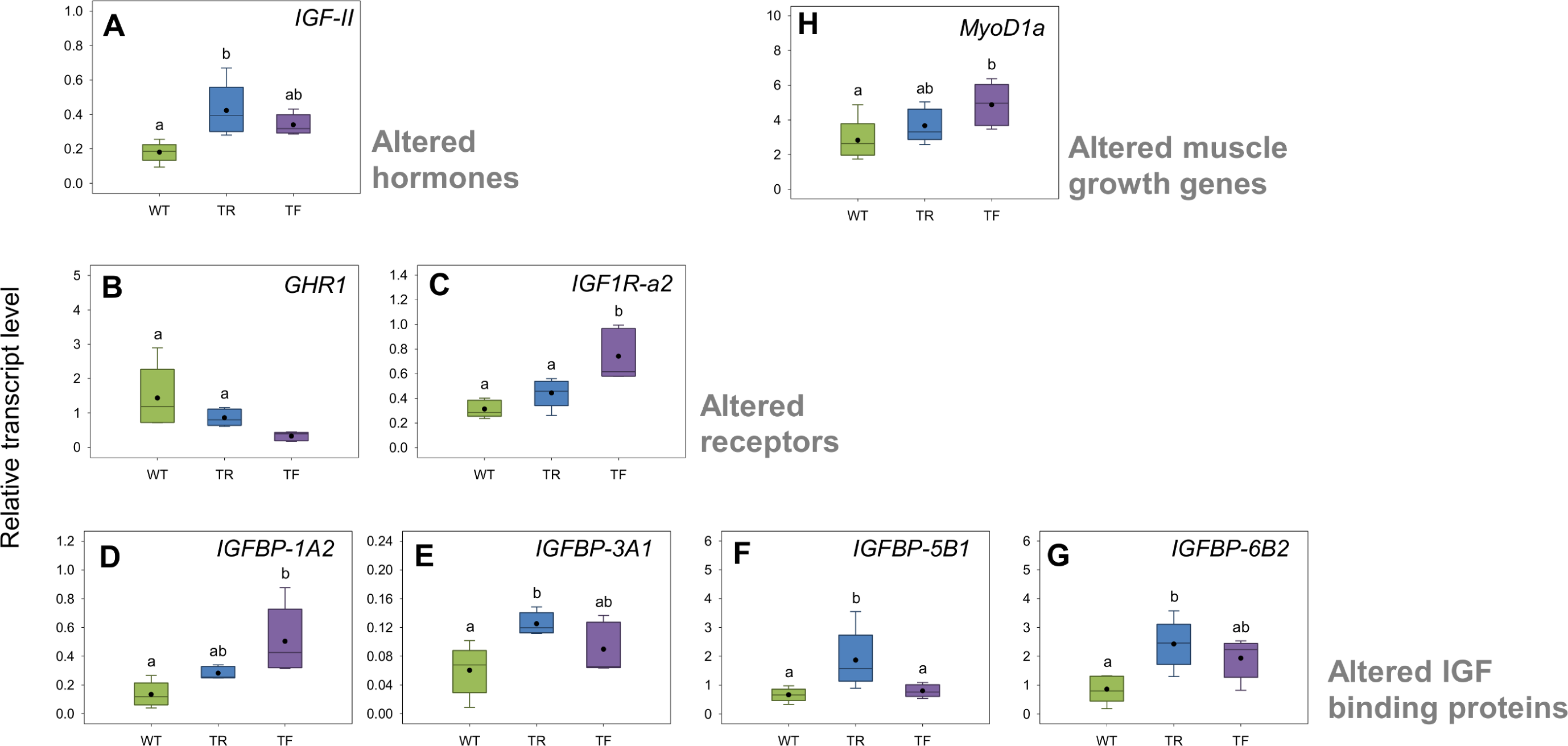
GH and IGF pathway genes with altered baseline expression comparing wild-type and GH transgenic fish. Box plots are shown for genes where a significant effect (*P*<0.05) was observed by one-way ANOVA comparing the three tested fish groups *(n=5* fish per group). Black circles within boxplots show mean transcript levels. Different letters indicate significant differences among fish groups (Tukey’s post hoc analysis). Full data (including means±s.d.) is given in Table 1.

**Table 1.** Differences in expression of GH and IGF system genes comparing wild-type to transgenic coho salmon groups

| Gene | P-value | WT transcript level | TR transcript level | TF transcript level |
| --- | --- | --- | --- | --- |
| Hormones |  |  |  |  |
| *GH* | 0.77 | 0.01±0.01 | 0.01±0.00 | 0.01±0.01 |
| *IGF-I* | 0.33 | 0.01±0.01 | 0.02 ±0.01 | 0.02±0.01 |
| *IGF-II* | 0.01 | 0.18±0.06 <sup>A</sup> | 0.42±0.15 <sup>B</sup> | 0.34±0.06 <sup>AB</sup> |
| Receptors |  |  |  |  |
| *GHR1* <sup>^</sup> | 0.001 | 1.43±0.90 <sup>A</sup> | 0.86±0.24 <sup>A</sup> | 0.32±0.13 <sup>B</sup> |
| *GHR2* | 0.24 | 0.73±0.67 | 0.56±0.13 | 0.43±0.09 |
| *IGF1R-a1* | 0.40 | 0.07±0.03 | 0.08±0.02 | 0.09±0.02 |
| *IGF1R-a2* | 0.001 | 0.31±0.07 <sup>A</sup> | 0.44±0.12 <sup>A</sup> | 0.74± 0.21 <sup>B</sup> |
| *IGF1R-b* | 0.36 | 0.07±0.02 | 0.10±0.03 | 0.09±0.04 |
| IGF binding proteins |  |  |  |  |
| *IGFBP-1A1* | 0.49 | 0.13±0.08 | 0.13±0.05 | 0.18±0.08 |
| *IGFBP-1A2* | 0.01 | 0.13±0.09 <sup>A</sup> | 0.28±0.04 <sup>AB</sup> | 0.50±0.23 <sup>B</sup> |
| *IGFBP-2A* <sup>^</sup> | 0.09 | 0.22±0.09 | 0.41±0.10 | 0.83±1.08 |
| *IGFBP-3A1* | 0.02 | 0.06±0.03 <sup>A</sup> | 0.13±0.02 <sup>B</sup> | 0.09±0.04 <sup>AB</sup> |
| *IGFBP-3A2* | 0.06 | 0.20±0.07 | 0.43±0.20 | 0.42±0.16 |
| *IGFBP-4* <sup>^</sup> | 0.69 | 1.03±1.30 | 0.91±0.42 | 0.65±0.14 |
| *IGFBP-5A* | 0.10 | 0.02±0.02 | 0.03±0.03 | 0.01±0.01 |
| *IGFBP-5B1* <sup>^</sup> | 0.01 | 0.66±0.23 <sup>A</sup> | 1.87±1.01 <sup>B</sup> | 0.80±0.22 <sup>A</sup> |
| *IGFBP-5B2* | 0.18 | 0.75±0.57 | 1.23±0.19 | 0.97±0.27 |
| *IGFBP-6B1* | 0.07 | 0.17±0.06 | 0.42±0.22 | 0.27±0.14 |
| *IGFBP-6B2* | 0.01 | 0.86±0.47 <sup>A</sup> | 2.42±0.83 <sup>B</sup> | 1.93±0.69 <sup>AB</sup> |
| Muscle growth status markers |  |  |  |  |
| *FBXO32* | 0.47 | 13.70±15.80 | 5.10±0.94 | 9.90±9.99 |
| *TNNI2* | 0.52 | 487.72±369.77 | 337.85±181.83 | 324.16±90.35 |
| *MYL1* | 0.89 | 230.07±136.61 | 198.63±75.88 | 221.97±92.12 |
| *MyoG* | 0.16 | 0.58±0.39 | 0.95±0.16 | 0.69±0.28 |
| *MyoD1a* | 0.04 | 2.83±1.21 <sup>A</sup> | 3.66±0.97 <sup>AB</sup> | 4.88±1.22 <sup>B</sup> |

Relative transcript levels (means±s.d, *n*=5) are shown as arbitrary units normalized to two empirically validated reference genes (*RpL13* and *ACTB*) and are quantitatively comparable across genes and among fish groups. Probability values are from one-way ANOVA testing for an effect of fish group. Different superscript letters highlight significant differences in transcript levels among groups according to Tukey's post-hoc test. The symbol "<sup>^</sup>" shows genes where the data required Box-Cox transformation.

### GH transgenesis alters skeletal muscle immune gene expression

To assess skeletal muscle responses to PGN, we measured transcript levels for markers of pro-inflammatory cytokines *(TNFα, IL-1β,* and *IL-8)* and acute phase proteins (SAA and *HAMP)* (Fig. 2A; Table S2) after Castro et al. 2015. A response to PGN was detected in each group, evidenced by a significant induction of all tested marker genes barring *TNFα* in TF (Fig. 2A; Table S2). The lack of *TNFα* response in TF was coupled to a respective 5.4 and 4.1-fold higher baseline expression vs. WT and TR (Fig. 2A; Table S2). In addition, the magnitude of observed responses of *TNFα, IL-1β* and *IL-8* was distinct among the fish groups; being highest in WT, intermediate in TR and lowest or non-existent in TF, leading to a significant treatment-x-group interaction (Fig. 2A; Table S2). In contrast, *SAA* and *HAMP* showed a highly significant induction in all groups, without any treatment-x-group interaction (Fig. 2A; Table S2).

**Figure 2.**
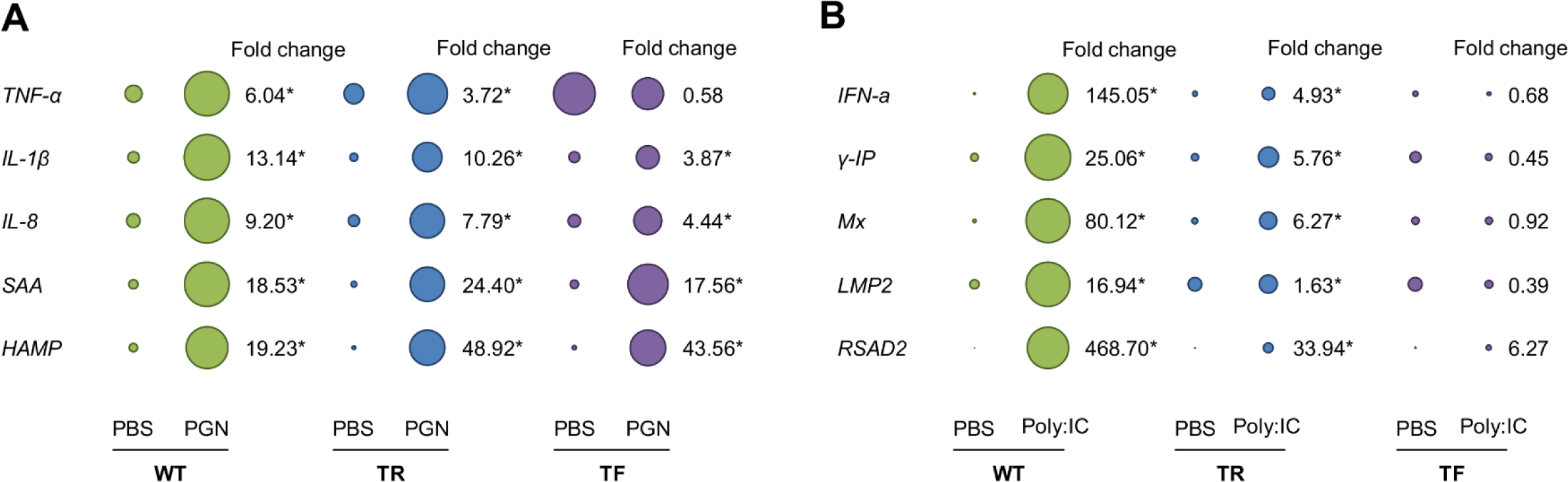
Impact of immune stimulation on the expression of host-defence genes comparing wild-type and GH transgenic fish. (A) PGN treatment. (B) Poly:IC treatment. Transcript expression is presented as bubble plots of sizes comparable among fish groups for each shown gene. The bubble area is proportional to mean transcript levels (n=5 fish, per group). Two-way ANOVA revealed that all the shown genes were significantly induced (P<0.05) by the immune stimulations (Table S2) and Tukey’s post hoc test identified significant changes in gene expression between control and treated fish, shown by asterisks next to accompanying fold-change values (calculated by dividing mean treatment by mean control, *n*=5 per mean). Full data (means±s.d.) is given in Table S2.

To assess skeletal muscle responses to Poly:IC, we measured transcript levels for markers of cytokines and proteins involved in the antiviral response *(IFN-a, y-IP, Mx, LMP2* and *RSAD2)* (Fig. 2B; Table S2) after Castro et al. 2015. Each tested marker was significantly induced in WT and TR, but not TF (Fig. 2B; Table S2). The responses of these genes showed a striking and consistent difference among groups, with between ~17 and 470 fold induction across the five genes in WT, compared to between ~1.6 and 34 fold in TR, and no upregulation in TF, leading to a highly significant treatment-x-group interaction (Fig. 2B; Table S2).

### GH transgenesis alters the responses of GH and IGF system genes to immune stimulation

PGN altered the expression of several GH and IGF pathway genes, with distinct responses observed among fish groups (Table 2). PGN had no effect on the expression of mRNAs encoding GH or IGF (Table 2). Considering the tested receptors, PGN had a significant overall effect on the expression of *GHR1, GHR2* and *IGF1R-b* (Table 2). These effects were different among the fish groups for GHR1, where a significant treatment-x-group interaction was observed (Table 2); GHR1 was significantly downregulated in WT by PGN, unchanged in TR, and significantly upregulated in TF by more than 5 fold (Table 2). The expression of several IGF-binding proteins was modified by PGN, often with distinct effects among groups (Table 2). *IGFBP-1A2* was significantly downregulated in TF, which followed a similar trend in TR, and an opposite trend in WT, where the same gene was upregulated (non-significant effect) (Table 2). Following a similar trend, *IGFBP-3A2* was significantly downregulated in both transgenic groups, while *IGFBP-6B2* was significantly downregulated in TR (Table 2). Considering the marker genes for muscle growth status, we observed no expression responses to PGN for genes encoding sarcomeric proteins along with *myoG*, while *MyoD1a* showed a strong and significant downregulation in TF specifically (Table 2).

**Table 2.** Effects of PGN on expression of GH and IGF system genes comparing wild-type to transgenic coho salmon groups

| Gene | P-value treatment | P-value treatment-x-group interaction | WT fold change; transcript level | TR fold change; transcript level | TR fold change; transcript level |
| --- | --- | --- | --- | --- | --- |
| Hormones |  |  |  |  |  |
| *GH* | 0.66 | 0.71 |  |  |  |
| *IGF-I* | 0.67 | 0.29 |  |  |  |
| *IGF-II* | 0.18 | <0.0001 |  |  |  |
| Receptors |  |  |  |  |  |
| *GHR1* ^ | 0.02 | <0.0001 | <u>0.33*</u> ; 0.47±0.21 <sup>A</sup> | <u>1.80</u> ; 1.55±0.46 <sup>B</sup> | <u>5.05*</u> ; 1.64±0.58 <sup>B</sup> |
| *GHR2* | 0.02 | - |  |  |  |
| *IGF1R-a1* | 0.19 | 0.002 |  |  |  |
| *IGF1R-a2* | 0.88 | <0.0001 |  |  |  |
| *IGF1R-b* | 0.03 | 0.09 |  |  |  |
| IGF binding proteins |  |  |  |  |  |
| *IGFBP-1A1* | 0.97 | <0.0001 |  |  |  |
| *IGFBP-1A2* | 0.01 | <0.0001 | <u>2.15</u> ; 0.29±0.11 <sup>A</sup> | <u>0.46</u> ; 0.13±0.05 <sup>A</sup> | <u>0.32*</u> ; 0.16±0.05 <sup>A</sup> |
| *IGFBP-2A* ^ | 0.22 | 0.001 |  |  |  |
| *IGFBP-3A1* | 0.27 | 0.78 |  |  |  |
| *IGFBP-3A2* | 0.01 | 0.001 | <u>1.77</u> ; 0.35±0.13 <sup>A</sup> | <u>0.33*</u> ; 0.14±0.02 <sup>A</sup> | <u>0.39*</u> ; 0.17±0.09 <sup>A</sup> |
| *IGFBP-4* | 0.05 | - |  |  |  |
| *IGFBP-5A* | 0.57 | 0.02 |  |  |  |
| *IGFBP-5B1* ^ | 0.90 | 0.09 |  |  |  |
| *IGFBP-5B2* | 0.58 | 0.47 |  |  |  |
| *IGFBP-6B1* | 0.87 | 0.001 |  |  |  |
| *IGFBP-6B2* | 0.05 | 0.002 | <u>1.89</u> ; 1.62±0.35 <sup>A</sup> | <u>0.51*</u> ; 1.24±0.52 <sup>A</sup> | <u>0.57</u> ; 1.10±0.38 <sup>A</sup> |
| Muscle growth status markers |  |  |  |  |  |
| *FBXO32* | 0.02 | 0.23 |  |  |  |
| *TNNI2* | 0.17 | 0.06 |  |  |  |
| *MYL1* | 0.11 | - |  |  |  |
| *MYOG* | 0.98 | 0.04 |  |  |  |
| *MyoD1a* | 0.002 | <0.0001 | <u>1.19</u> ; 3.38±0.87 <sup>A</sup> | <u>0.82</u> ; 3.00±0.65 <sup>AB</sup> | <u>0.30*</u> ; 1.46±0.33 <sup>B</sup> |

Fold change values (underlined: calculated by dividing the mean control and mean PGN treatment transcript levels, $n=5$ fish per mean) are given for genes where a significant effect of treatment and a significant treatment-x-group interaction was recorded by two-way ANOVA (the symbol '\*' indicates a significant expression change according to Tukeys post-hoc test). For the same genes, relative transcript levels are shown post-PGN treatment (means±s.d, $n=5$ ) and represent arbitrary units normalized to two empirically validated reference genes (*Rpl13* and *ACTB*) that are quantitatively comparable across genes and among fish groups (different superscript letters highlight significant differences among groups according to Tukeys post-hoc test). The symbol '^' shows genes where the data required Box-Cox transformation. The symbol '-' shows genes where a Kruskal-Wallis test was done testing for a treatment effect only.

Poly:IC had a major effect on the expression of GH and IGF pathway genes, with many showing distinct responses among wild-type and GH transgenic fish, reflected in significant treatment-x-group interaction effects (Table 3). Among the tested hormones, *IGF-II* showed a significant downregulation in both transgenic groups (Table 3). For the tested receptors, *GHR1* was significantly induced in TF, while *IGF1R-b* showed a significant downregulation in both transgenic groups (Table 3). Several IGFBP mRNAs were significantly altered by Poly:IC, with *IGFBP-1A2* showing significant upregulation in WT, but significant downregulation in both transgenic groups (Table 3). A similar pattern was observed for *IGFBP-2A, IGFBP-5B2, IGFBP-6B1* and *IGFBP-6B2*, with significant downregulation in both transgenic groups (Table 3). Considering the markers of muscle growth status, *MyoD1a* was significantly downregulated by Poly:IC in TF (Table 3).

**Table 3.** Effect of Poly:IC on expression of GH and IGF system genes comparing wild-type to transgenic coho salmon groups

| Gene | P-value treatment | P-value treatment-x-group interaction | WT fold change; transcript level | TR fold change; transcript level | TR fold change; transcript level |
| --- | --- | --- | --- | --- | --- |
| Hormones |  |  |  |  |  |
| *GH* | 0.48 | 0.63 |  |  |  |
| *IGF-I* | 0.74 | 0.15 |  |  |  |
| *IGF-II* ^ | <0.0001 | 0.001 | <u>1.20</u> ; 0.22±0.03 <sup>A</sup> | <u>0.43*</u> ; 0.18±0.05 <sup>A</sup> | <u>0.46*</u> ; 0.15±0.08 <sup>A</sup> |
| Receptors |  |  |  |  |  |
| *GHR1* ^ | 0.05 | <0.0001 | <u>0.44</u> ; 0.64±0.38 <sup>A</sup> | <u>1.34</u> ; 1.15±0.14 <sup>A</sup> | <u>3.80*</u> ; 1.24±0.30 <sup>A</sup> |
| *GHR2* ^ | 0.33 | 0.38 |  |  |  |
| *IGF1R-a1* | 0.56 | 0.06 |  |  |  |
| *IGF1R-a2* | 0.17 | <0.0001 |  |  |  |
| *IGF1R-b* | 0.03 | <0.0001 | <u>1.70</u> ; 0.12±0.04 <sup>A</sup> | <u>0.34*</u> ; 0.03±0.01 <sup>B</sup> | <u>0.32*</u> ; 0.03±0.01 <sup>B</sup> |
| IGF binding proteins |  |  |  |  |  |
| *IGFBP-1A1* | 0.07 | 0.02 |  |  |  |
| *IGFBP-1A2* ^ | 0.05 | <0.0001 | <u>2.52*</u> ; 0.34±0.10 <sup>A</sup> | <u>0.43*</u> ; 0.12±0.04 <sup>B</sup> | <u>0.30*</u> ; 0.15±0.04 <sup>B</sup> |
| *IGFBP-2A* ^ | 0.05 | 0.001 | <u>2.91</u> ; 0.64±0.73 <sup>A</sup> | <u>0.46*</u> ; 0.19±0.06 <sup>A</sup> | <u>0.24*</u> ; 0.20±0.13 <sup>A</sup> |
| *IGFBP-3A1* | 0.02 | 0.14 |  |  |  |
| *IGFBP-3A2* | 0.25 | 0.001 |  |  |  |
| *IGFBP-4* ^ | 0.70 | 0.61 |  |  |  |
| *IGFBP-5A* ^ | 0.14 | 0.29 |  |  |  |
| *IGFBP-5B1* ^ | 0.70 | 0.11 |  |  |  |
| *IGFBP-5B2* | 0.01 | 0.03 | <u>1.16</u> ; 0.87±0.22 <sup>A</sup> | <u>0.50*</u> ; 0.61±0.08 <sup>A</sup> | <u>0.54*</u> ; 0.52±0.26 <sup>A</sup> |
| *IGFBP-6B1* ^ | <0.0001 | 0.01 | <u>0.90</u> ; 0.15±0.05 <sup>A</sup> | <u>0.20*</u> ; 0.09±0.04 <sup>A</sup> | <u>0.33*</u> ; 0.09±0.06 <sup>A</sup> |
| *IGFBP-6B2* | 0.01 | 0.01 | <u>1.44</u> ; 1.24±0.33 <sup>A</sup> | <u>0.54*</u> ; 1.30±0.54 <sup>A</sup> | <u>0.42*</u> ; 0.81±0.32 <sup>A</sup> |
| Muscle growth status markers |  |  |  |  |  |
| *FBXO32* ^ | 0.20 | 0.13 |  |  |  |
| *TNNI2* | 0.20 | 0.03 |  |  |  |
| *MYL1* | 0.33 | 0.72 |  |  |  |
| *MYOG* | 0.71 | 0.04 |  |  |  |
| *MyoD1a* | <0.0001 | 0.01 | <u>0.83</u> ; 2.34±0.57 <sup>A</sup> | <u>0.60</u> ; 2.19±0.19 <sup>A</sup> | <u>0.37*</u> ; 1.79±0.43 <sup>A</sup> |

Fold change values (underlined: calculated by dividing the mean control and mean Poly:IC treatment transcript levels, $n=5$ fish per mean) are given for genes where a significant effect of treatment and a significant treatment-x-group interaction was recorded by two-way ANOVA (the symbol '\*' indicates a significant expression change according to Tukeys post-hoc test). For the same genes, relative transcript levels are shown post-Poly:IC treatment (means±s.d, $n=5$ ) and represent arbitrary units normalized to two empirically validated reference genes (*RpL13* and *ACTB*) that are quantitatively comparable across genes and among fish groups (different superscript letters highlight significant differences among groups according to Tukeys post-hoc test). The symbol '^' shows genes where the data required Box-Cox transformation. The symbol '-' shows genes where a Kruskal-Wallis test was done testing for a treatment effect only.

## DISCUSSION

Animals have evolved complex defence mechanisms to eradicate invading pathogens, which are energetically expensive (e.g. Bonneaud et al. 2003; Lochmiller and Deerenberg, 2000; Cutrera et al. 2010). Recent studies in fish have revealed that the expression of certain genes from the IGF system is altered during the host defence response to bacterial and viral infections, consistent with a downregulation of growth and reallocation of resources towards immune function (Pooley et al., 2013; Alzaid et al., 2016a, 2016b). Until now, little consideration has been given to the regulation of growth pathways following immune stimulation in skeletal muscle, which is the most important target tissue for energetic investment and storage. Here, we provide evidence for cross-talk between growth and immune function in coho salmon muscle, which is disrupted by GH transgenesis as well as its impacts on growth rate and physiological status.

### Altered expression of GH and IGF system genes by GH transgenesis

Several GH and IGF system genes, along with muscle growth genes, showed altered expression in GH transgenic coho, often with responses differing among TR and TF groups. We observed a significant elevation of *IGF-II* and *IGFBP-5B1* expression in TR, but not TF, compared to WT. A past study on the same type of coho groups documented a life-long reduction in muscle fibre production rate in TR when compared to both TF and WT, accompanied by downregulation of genes involved in myotube formation (Johnston et al. 2014). This observation can be explained by energetic savings linked to ionic homeostasis in muscle (the ‘optimal fibre size’ hypothesis; see Johnston et al. 2012), allowing the TR group to allocate additional resources into foraging, demanded by their increased appetite (Johnston et al. 2014). IGFBP-5 is known to promote muscle differentiation by binding IGF-II, and switching on an auto-regulatory loop that induces further *IGF-II* expression (Ren et al., 2008). The concurrent upregulation of *IGF-II* and *IGFBP-5B1* in TR specifically may be connected to this auto-regulatory loop, and moreover related to the reduced rate of muscle fibre production, potentially driving muscle growth towards hypertrophy at the expense of muscle fibre production (Johnston et al. 2011).

Other molecular markers of myogenesis were altered by GH transgenesis in the TF group specifically, notably an increase in the expression of *MyoD1a,* which encodes a salmonid duplicate for a transcription factor from the myogenic regulatory factor family, which also includes MyoG (Macqueen et al. 2007). This finding points towards differences in myogenic development influenced by growth rate and nutritional status in GH transgenics. A past study showed that *MyoD1a* and *MyoG* were co-expressed during development of Atlantic salmon primary muscle cell cultures, each peaking in expression during myogenic differentiation (Bower and Johnston, 2010). However, in our study, *MyoG* was not changed by GH transgenesis, in contrast to past reports in coho salmon (Overturf et al. 2010), suggesting differences in GH-dependent regulation of *MyoD1a* and *MyoG* that may relate to unique functions in myogenesis. We also noted a lack of regulation for mRNAs encoding FBXO32 (as reported in Overturf et al. 2010) and muscle structural proteins in both GH transgenic groups, suggesting a limited impact of GH on the turnover of muscle structural proteins, even for the rapidly growing TF group.

We also observed low *GH* transcript levels in all the fish groups, consistent with past data for skeletal muscle (Raven et al., 2008; Devlin et al., 2009; Johnston et al., 2014). However, *GH* expression was reported elsewhere to increase in skeletal muscle of GH transgenics (Raven et al., 2008; Devlin et al., 2009; Johnston et al., 2014), which was not observed here. *GHR1* expression was highly reduced in the TF group, which could represent a compensatory feedback response to the ubiquitous presence of circulating GH available to muscle cells. Among three distinct *IGF1R* genes retained in salmonids (Alzaid et al. 2016a), only *IGF1R-a2* showed a response to GH transgenesis, being upregulated in the TF group, a reciprocal pattern to *GHR1*. Considering a recent study that demonstrated an upregulation of *IGF1R* in muscle cell cultures treated with IGF-I (Huang et al. 2011), this finding may relate to the elevated level of IGF-I in serum of the TF group (Raven et al., 2008). The lack of upregulation of *IGF1R-a1* in the same context provides clear evidence for divergent regulation since the salmonid-specific genome duplication event.

The skeletal muscle expression of several IGFBPs in addition to *IGFBP-5B1* (discussed above) was altered by GH transgenesis, again with differences among TF and TR groups. This suggests that the interaction of IGFs with IGF-1R is regulated by a unique prolife of IGFBPs depending on growth rate, in addition to GH transgenesis. For example, *IGFBP-1A2* was significantly higher in TF than WT, while *IGFBP-3A1* and *IGFBP-6B2* were significantly higher in TR than WT. IGFBP-1 is classically considered a negative regulator of IGF signalling that sequesters IGF-I away from IGF-1R, and is upregulated under catabolic conditions to limit growth (e.g. Lang et al. 2003; Kajimura et al. 2005; Shimizu and Dickhoff, 2017), though its auto/paracrine role in muscle remains poorly understood. The increased expression of *IGFBP-1A2* in the TF group is thus difficult to explain in terms of its classic function, but this could be related to IGF-independent roles (Wheatcroft, and Kearney, 2009). The upregulation of *IGFBP-6B2* specifically in TR is notable, as IGFBP-6 subtypes are thought to bind to IGF-II with higher affinity than IGF-I (Bach, 2016) and this expression profile is coupled to the aforementioned increase in *IGF-II* in the same group, suggesting a potential functional relationship.

### GH transgenesis disrupts immune status of skeletal muscle

We observed differences in the skeletal muscle responses of immune genes among the three coho groups following injections with mimics of bacterial (PGN) and viral (Poly:IC) infection. In response to PGN, the pro-inflammatory cytokines *IL-1β, IL-8* and *TNFα* were strongly induced in WT, confirming the anticipated inflammatory response. While TR and TF groups showed a comparable response, the magnitude of cytokine induction was lower than WT, and particularly attenuated in TF, which failed to upregulate *TNF-α* (Fig. 2A, Table S2). These findings suggest that the GH transgenic fish have a reduced inflammatory response, particularly when maximal growth rate is being achieved. However, genes encoding the acute phase proteins Serum amyloid A and Hepcidin were induced at comparable levels in all coho groups, indicating that GH transgenesis does not compromise all effectors of innate immunity in skeletal muscle.

The altered expression of pro-inflammatory cytokines observed in GH transgenics is indicative of a disruption in normal cross-talk between skeletal muscle and immune function (reviewed in Pillon et al. 2013). This could relate to differences in the number of immune cells resident within the muscle, for example macrophages and granulocytes, given past reports that GH transgenic salmon develop fewer white cells than wild-type fish (Kim et al., 2013). The constitutive up-regulation of *TNF-α* in the TF group, suggests that growth rate and/or nutritional status modulate the impact of GH on *TNF-α* regulation, and imply a permanent inflammatory state, which classically would be deemed pathological (Pillon et al. 2013). On the other hand, TNF-α, along with other pro-inflammatory cytokines such as IL-1, positively regulate muscle development (e.g. Chen et al. 2007; Costamagna et al. 2015). Hence, the disrupted cytokine expression has implications for myogenesis in addition to health status. GH transgenic coho were previously shown to have higher susceptibility to the bacterium *Aeromonas salmonicida* (Kim et al., 2013), which implies a systematic attenuation of innate immunity. However, more data will be needed to confirm this hypothesis at the molecular level, including for primary immune tissues, as the present immune expression data was restricted to skeletal muscle.

In response to Poly:IC, the expression levels of all selected markers of the antiviral response showed a robust induction in skeletal muscle of WT. However, the same genes showed a highly reduced up-regulation in TR and remarkably, were not induced in the TF group. The selected marker genes for the antiviral response are effectors of Type-I interferon signalling (e.g. Martin et al. 2007), which acts to control viral replication on several levels, including by inhibition of host translation and protein synthesis (Haller et al. 2007; Sadler and Williams, 2008). With that in mind, our results can potentially be explained by the presence of ubiquitous signalling towards new protein synthesis in the TF group, likely through the IGF-PI3K-Akt-mTOR pathway downstream of GH (Egerman and Glass, 2014). Past studies have provided evidence for cross-talk between interferon signalling and the IGF-PI3K-Akt-mTOR pathway (e.g. Kaur et al. 2008), providing possible mechanisms whereby the antiviral response could be downregulated in favour of protein synthesis. However, as gene markers for Type-I interferon signalling showed attenuated expression in the TR group, which have wild-type growth rate and IGF-1 levels (Raven et al. 2008), our findings also imply a direct role for GH in limiting expression of these antiviral genes. It will be important in the future to establish whether the absence of antiviral gene induction in muscle of GH transgenic salmon is a consequence of systematic attenuation in antiviral immunity, or a tissue-specific effect caused by muscle’s key role as a target for growth and protein synthesis.

### GH transgenesis alters cross-talk between the GH/IGF system and immune function

Several GH and IGF system genes showed altered regulation to immune stimulation in GH transgenics. This effect was particular evident in response to Poly:IC, mirroring the strong attenuation of the antiviral response, and supporting the hypothesis that altered immune function in transgenics is directly influencing growth gene expression and ‘normal’ cross-talk between immunity and growth.

However, the observed changes in growth gene expression are complex, and not straightforward to interpret in full. It is also important to remember the baseline levels of gene expression in the different coho groups, when interpreting observed changes in gene regulation. For example, *IGF-II* was significantly downregulated by Poly:IC in both transgenic groups, but not WT, yet there were no differences in the post-treatment transcript levels comparing any group - hence, *IGF-II* downregulation reduced expression to the wild-type level. As *IGF-II* promotes muscle growth (e.g. Ren et al., 2008), this provides evidence that GH transgenics may be suppressing growth signalling to the wild-type level during a viral immune response. However, as *IGF-II* downregulation was equally evident in both TR and TF groups, the underlying mechanisms must result from GH transgenesis *per se,* which is less consistent with a scenario where the fast-growing TF group is reallocating available resources to immune function. In a similar respect, mRNAs encoding several IGFBP sub-types *(IGFBP-6B2* for both treatments; *IGFBP-6B1* for Poly:IC; *IGFBP-5B2* for Poly:IC and *IGFBP-2A* for Poly:IC) showed significant differences in regulation across the fish groups, but had similar mRNA expression levels across all groups after immune mimic treatment. Invariably, this was coupled to downregulation of IGFBP expression in both the GH transgenic groups, suggesting these altered responses to immune stimulation were caused by GH transgenesis, rather than its impact on growth rate. The affected IGFBP subtypes are functionally diverse, and have previously been accredited with both positive (e.g. IGFBP-3: Foulstone et al. 2003; IGFBP-5: Ren et al. 2008) and inhibitory (e.g. IGFBP-2: Swiderski et al. 2016; IGFBP-6: Pooley et al. 2013) impacts on muscle growth. Hence, it is difficult to predict the effects that downregulation of multiple IGFBP subtypes will have in the GH transgenics, both in terms of muscle growth, but also several other metabolic phenotypes known to be regulated by this protein family (Jones and Clemmons, 1995).

In other cases, patterns of gene expression in GH transgenics were consistent with an upregulation of growth signalling in response to immune challenge, when wild-type fish were downregulating growth. In response to both PGN and Poly:IC, the GH receptor *GHR1* was strongly upregulated in TF, and the post-treatment level of *GHR1* mRNA in GH transgenics was higher than for the WT group, which downregulated *GHR1* (significant effect for PGN only) in response to immune stimulation. As GH receptors are required for GH signalling, these patterns suggest that regulatory mechanisms present in wild-type fish that restrict growth during an immune challenge are altered in GH transgenics when maximal growth is being achieved. Wild-type fish also showed an upregulation of *IGFBP-1A2* in response to immune stimulation (significant for Poly:IC), which may serve to restrict IGF signalling by sequestering IGF hormones away from IGF-1R. In GH transgenics, the opposite trend was observed, with downregulation of *IGFBP-1A2* mRNA to a level below that of wild-type, which might be expected to comparatively promote growth by increasing IGF availability to IGF-1R.

We also observed a strong downregulation of *IGF1R-b* for both GH transgenic groups in response to Poly:IC specifically. As this IGF receptor subtype was expressed at the same levels in the unstimulated fish, this resulted in both GH transgenic groups being expressed at much lower mRNA levels post-immune stimulation. IGF receptors are important for muscle development (Mavalli et al. 2010) and the IGF-1Rb subtype of zebrafish is known to have evolved a specific function in regulating motor neuron development (Schlueter et al. 2006). Hence the downregulation of *IGF1R-b* in GH transgenics in response to Poly:IC has implications for both muscle development and function.

Much future work is needed to better understand the complex role played by GH transgenesis in regulating the responses of growth systems to immune challenge in teleosts, which should involve consideration of additional tissues, as well as a deeper dissection of molecular signatures using global transcriptomic or proteomic approaches.

### Broader perspectives

This study demonstrates that GH transgenesis, and the associated enhancement in growth rate, disrupts immune function in coho salmon muscle, with downstream impacts on cross-talk between immunity and growth systems. It will be important to further explore whether the effects of rapid growth rate induced during domestication and/or by selection - occurring over multiple generations with opportunity for co-adaptation and compensatory selective responses in interacting pathways - are the same or different from GH transgenesis. While many parallels in endocrinology and gene expression are observed in domesticated and GH transgenic fish relative to wild-type, differences have also been detected in multiple physiological pathways (Fleming et al. 2002; Tymchuck et al. 2009; Devlin et al. 2009; Devlin et al. 2013). Taken with past reports of altered immune function in GH transgenics (Jhingan et al. 2003; Kim et al. 2013), our finding that the muscle of GH transgenic salmon achieving maximal growth lacked any detectable antiviral response, and showed a baseline upregulation of *TNF-α* expression, has implications for the health/welfare of GH transgenic strains destined for aquaculture, where rapid growth is the central goal of production. Finally, our data have ramifications for environmental risk assessments (Devlin et al. 2015) tasked with determining the fitness and potential impacts of genetically engineered fishes should they enter natural environments.

## Acknowledgements

We thank Dwight Causey (Macqueen lab, University of Aberdeen) for providing critical comments on the manuscript.

## Author Contributions

Designed research: DJM, RHD, JHK, SAMM; performed animal experiments and sampling: JHK, RHD; performed molecular biology and qPCR analysis: AA; analysed qPCR data: AA, DJM; performed statistics: AA, DJM. The manuscript, including tables and figures, was drafted by AA and DJM and finalized with extensive inputs from all authors.

---

**Table S1.**
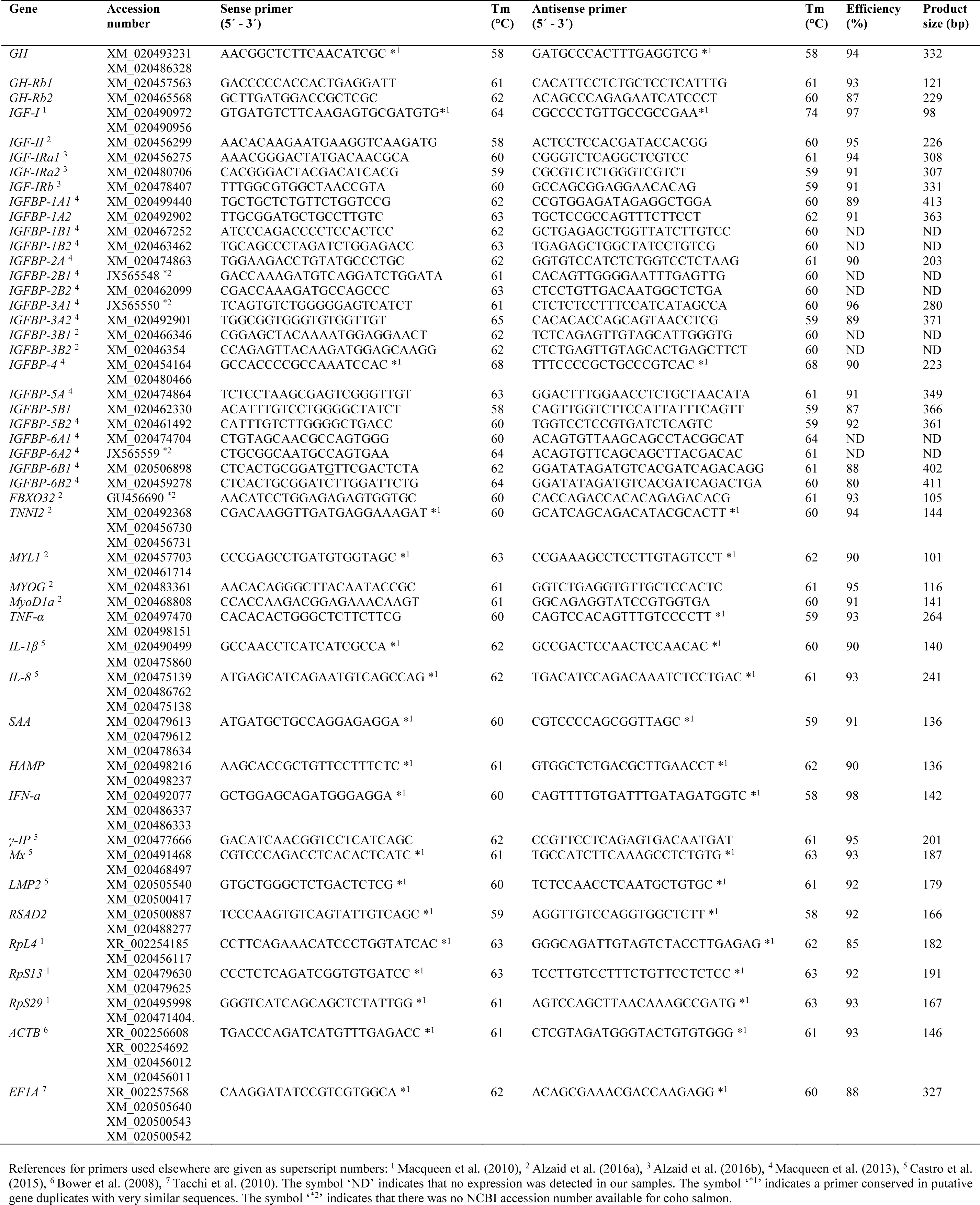
Details of primers used for qPCR analyses

**Table S2.** Effects of immune stimulation on expression of host defence genes comparing wild-type to transgenic coho salmon groups

| Treatment | Gene | P-value<br>treatment | P-value<br>treatment-x-group<br>interaction | WT transcript level |  | TR transcript level |  | TF transcript level |  |
| --- | --- | --- | --- | --- | --- | --- | --- | --- | --- |
|  |  |  |  | Control | Treatment | Control | Treatment | Control | Treatment |
| PGN | *TNF-α* | <0.0001 | 0.001 | 11.40±6.23 | 68.88±25.69 | 14.88±4.16 | 55.33±18.67 | 61.67±12.93 | 36.00±14.31 |
| | *IL-1β* | <0.0001 | <0.0001 | 5.42±6.68 | 71.22±23.39 | 3.02±0.86 | 30.98±8.24 | 5.03±1.98 | 19.46±10.04 |
| | *IL-8* ^ | <0.0001 | 0.01 | 7.48±6.76 | 68.81±19.18 | 5.32±0.46 | 41.49±5.99 | 6.43±1.35 | 28.54±15.67 |
| | *SAA* ^ | <0.0001 | 0.44 | 3.71±5.99 | 68.66±22.92 | 1.72±1.04 | 41.86±6.68 | 3.21±1.83 | 56.42±24.22 |
| | *HAMP* | <0.0001 | - | 3.17±5.62 | 60.87±24.97 | 0.88±0.41 | 43.17±7.64 | 1.04±0.86 | 45.14±20.09 |
| Poly:IC | *IFN-α* ^ | <0.0001 | <0.0001 | 0.37±0.32 | 54.30±27.60 | 1.26±0.21 | 6.23±4.03 | 1.34±0.63 | 0.91±0.26 |
| | *γ-IP* ^ | 0.001 | <0.0001 | 2.77±1.45 | 69.44±21.49 | 2.56±0.79 | 14.75±10.93 | 4.96±3.49 | 2.23±0.69 |
| | *Mx* ^ | <0.0001 | <0.0001 | 0.82±0.71 | 65.41±23.45 | 1.77±1.13 | 11.12±5.98 | 2.60±1.10 | 2.40±2.10 |
| | *LMP2* ^ | 0.001 | <0.0001 | 3.90±3.23 | 66.06±28.56 | 7.41±3.85 | 12.10±7.67 | 7.40±4.38 | 2.85±0.25 |
| | *RSAD2* ^ | <0.0001 | 0.002 | 0.12±0.06 | 57.97±30.65 | 0.12±0.08 | 4.02±2.70 | 0.23±0.16 | 1.46±2.00 |

Relative transcript levels (means ± s.d, $n=5$ ) are shown as arbitrary units normalized to two empirically validated reference genes (*RpL13* and *ACTB*) and are quantitatively comparable across genes and among fish groups. Probability values shown are from a two-way ANOVA analysis. The symbol '^' shows genes where the data required Box-Cox transformation. The symbol '-' shows genes where a Kruskal-Wallis test was done testing for a treatment effect only.

